# From enhancers to genome conformation: complex transcriptional control underlies expression of a single herpesviral gene

**DOI:** 10.1101/2023.07.08.548212

**Authors:** David W Morgens, Leah Gulyas, Alejandro Rivera-Madera, Annabelle S Souza, Britt A Glaunsinger

## Abstract

Complex transcriptional control is a conserved feature of both eukaryotes and the viruses that infect them. Here, we illustrate this by combining high-density functional genomics, expression profiling, and viral-specific chromosome conformation capture to define with unprecedented detail the transcriptional regulation of a single gene, ORF68, from Kaposi’s sarcoma-associated herpesvirus (KSHV). We first identified seven cis-regulatory regions by densely tiling the ∼154 kb KSHV genome with CRISPRi. A parallel Cas9 nuclease screen indicated that three of these regions act as promoters of genes that regulate ORF68. RNA expression profiling demonstrated that three more of these regions act by either repressing or enhancing other distal viral genes involved in ORF68 transcriptional regulation. Finally, we tracked how the 3D structure of the viral genome changes during its lifecycle, revealing that these enhancing regulatory elements are physically closer to their targets when active, and that disrupting some elements caused large- scale changes to the 3D genome. These data enable us to construct a complete model revealing that the mechanistic diversity of this essential regulatory circuit matches that of human genes.

## Introduction

Underlying the lifecycle of all DNA viruses is a highly regulated cascade of viral gene transcription. In human herpesviruses, many of these genes are transcribed in a host-like manner; nuclear dsDNA viral genomes can be chromatinized by human histones,^1^ decorated with enhancer marks,^2^ driven by human transcription factor binding,^3^ and even form transcription-associated domains with human CTCF and cohesion.^4,5^ Exhaustively identifying and characterizing these regulatory features can both help us understand the biology of human pathogens such as Kaposi sarcoma-associated herpesvirus (KHSV) – a major cause of cancer in AIDS and other immunocompromised patients – and also contribute to our knowledge of human transcriptional regulation.

Numerous examples of non-coding regulatory sequences have been found in DNA viruses. Enhancer sequences include the first described enhancer on SV40,^6^ the EIIA enhancer on adenovirus,^7^ the major immediate enhancer element in the betaherpesvirus human cytomegalovirus,^8,9^ and a recent proposal that the terminal repeats of KSHV act as enhancers.^10^ Other viral regulatory elements act through expression of non-coding elements. In murine gammaherpesvirus 68 there are tRNA-like elements that control latency and egress.^11,12^ In KSHV, many different functional elements beyond coding mRNAs are transcribed, including miRNAs that regulate cancer phenotypes,^13,14^ origin RNAs that regulate viral DNA replication,^15,16^ circular RNAs,^17,18^ and long ncRNAs with various functions.^19–23^ While these individual events have been explored to different degrees, systematic searches for regulatory sequences have been limited by traditional methods that rely on deletions to perturb noncoding elements, which in the densely encoded KSHV genome may have unintended effects on surrounding elements.

For the human genome, the discovery of functional regulatory sequences has been greatly accelerated by the use of CRISPR interference or CRISPRi.^24–28^ Notably, by recruiting repressive chromatin regulators to DNA, CRISPRi can repress gene expression not only through proximal promoter elements but also by perturbing distal, enhancer elements. While this tool has been applied widely on the host, it can also effectively repress transcription from the KSHV genome,^29^ and thus has the potential to provide deep insight into the components and structure of viral transcriptional networks. While searches for human regulatory regions are limited to predicted enhancers or nearby regions, the viral regulatory regions will be necessarily contained within the relatively compact viral genome and thus be amenable to exhaustive characterization.

Here we combine CRISPRi with a library of guide RNAs densely tiling the KSHV genome, allowing a thorough interrogation of potential regulatory activity controlling the expression of a single viral gene, ORF68. ORF68 was selected as a proof of concept for this study as it has a simple early- expression pattern^30,31^ but plays an essential role late in the viral life cycle during packaging of new viral DNA.^32,33^ By complementing CRISPRi with a Cas9 nuclease screen and transcriptional profiling, we identified promoters that control expression through their associated coding regions, as well as non-coding regulatory elements that comprise a surprisingly sophisticated network to regulate ORF68 transcription. Finally, we use viral-specific chromosome conformation capture to measure physical interactions between regulatory regions and their distal targets and demonstrate how disruption of these regions changed the 3D structure of the viral genome. These findings illustrate the power of this approach for mapping viral gene regulatory networks on a genome-wide scale.

## Results

### CRISPRi tiling identifies regulatory regions across the viral genome

We sought to define the layers of regulation underlying the expression of an individual viral gene, using the nuclear replicating dsDNA virus KSHV. We selected KSHV ORF68 as our model gene, as it is required for progeny virion production, has no known direct transcriptional regulators, and its expression initiates early in the lytic cycle and stays on throughout the rest of the viral lifecycle^30,31^. KSHV gene regulation can be readily studied using the renal carcinoma cell line iSLK, a well-established model for the KSHV lifecycle that includes a doxycycline-inducible version of the KSHV lytic transactivator ORF50 to facilitate lytic reactivation from latency.

We began construction of this regulatory network by querying how silencing of each KSHV locus using CRISPRi influenced ORF68 expression. We latently infected iSLK cells with a version of KSHV containing a HaloTag fused to the N-terminus of ORF68 at the endogenous locus (HaloTag- ORF68),^34^ allowing us to directly measure ORF68 protein levels, as well as a constitutively expressed GFP reporter that marks infected cells. Additionally, we lentivirally introduced a constitutive dCas9-KRAB fusion (CRISPRi), which would be recruited to a targeted viral region upon delivery of a sgRNA. We then delivered a library of guide RNAs densely tiling the KSHV genome with an average of one guide every 8 base pairs.^34^ After four days, the virus was reactivated, and cells were treated with a fluorescent HaloTag ligand to monitor ORF68 protein levels. 24 hours post-reactivation, cells were fixed and sorted for high and low ORF68 protein expression (**Figure 1a**). By sequencing the sgRNA locus from both populations, we calculated an average guide enrichment from two replicates. A negative value signifies that the guide was enriched in the low ORF68 signal population, indicating that silencing that locus inhibits the expression of ORF68. Similarly, a positive value signifies that the guide was enriched in the high ORF68 signal population, indicating that silencing that locus increases expression of ORF68 (**Figure S1a,b**).

**Figure 1.**
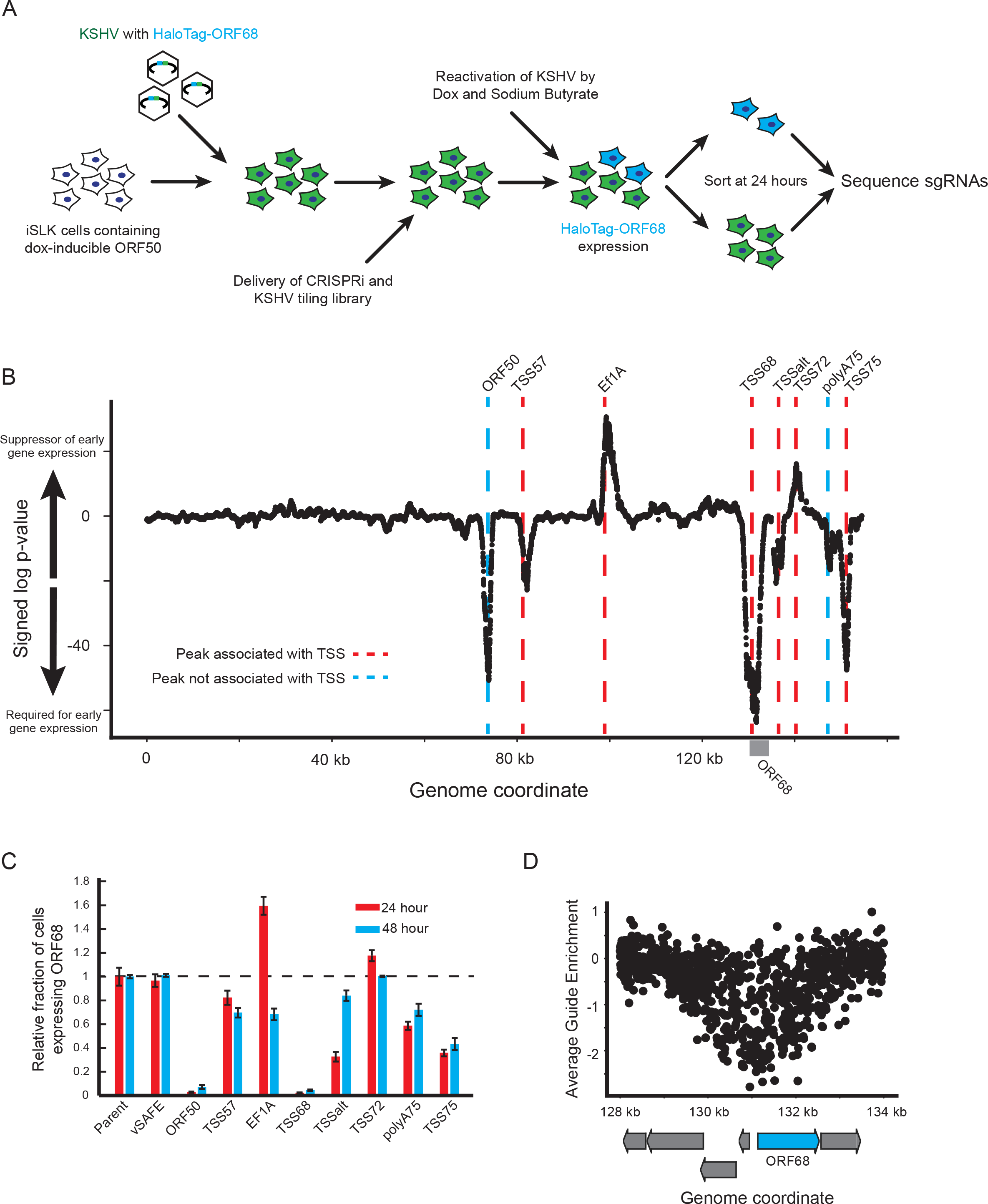
*CRISPRi screen identifies novel viral regulatory regions*. A) Schematic of screen. The viral genome encodes a constitutive fluorescent marker (green) and a HaloTag-ORF68 fusion (blue). B) Summary of results from the CRISPRi screen. X-axis identifies the genome coordinate on the BAC16 KSHV genome. y-axis represents the log-transformed p-value of each locus relative to negative control distribution. Red dotted lines indicate the location of transcriptional start sites. Blue dotted line identifies the two peaks that do not associate with a TSS. C) Validation of pooled guides targeting each peak. Three guides were used to target each locus identified on the x-axis. Y-axis displays the relative percent of cell expressing the HaloTag-ORF68 for the 24 or 48 hour post lytic reactivation time points. Error bars represent standard error of four replicates from independent reactivations. D) Enrichment of individual guides at the ORF68 locus. Each dot represents a single guide, with the target location displayed on the x-axis and the average enrichment from two replicates on the y-axis. Arrows represent coding regions of ORF68 (in blue) and surrounding genes in grey.

We performed a sliding window analysis and identified 8 loci with target guide RNA scores that differed significantly from negative controls (**Figure 1b; Supplementary Data 1, 2**). Our strongest signal corresponded to a peak poised immediately upstream of HaloTag-ORF68 itself, confirming successful transcriptional inhibition of the locus by CRISPRi. We observed that the center of most other peaks also corresponded with known transcriptional start sites (TSSs); since we expect CRISPRi to work primarily by impeding transcription, these peaks are named by their nearest TSS for simplicity (**Figure S1c-h**). For example, we find one peak near the TSS of ALT, a lncRNA of unknown function which runs antisense to many genes expressed during latency,^22^ that we will refer to as TSSalt (**Figure S1e**). The exceptions include one peak that roughly maps to the ORF50 coding locus (**Figure S1c**) – likely targeting the exogenous copy of ORF50 used to reactivate the virus – as well as an additional peak which does not correspond to a previously described TSS (**Figure S1h**)^35^ but is located near the shared polyA sites of ORF75, ORF74, K14, K15, and the ALT lncRNA.^22,36^ We will refer to these as ORF50 and polyA75 respectively.

We next validated these 8 regions’ effect on ORF68 by delivering a pool of three CRISPRi guides per locus and measuring the percentage of cells expressing HaloTag-ORF68 at two timepoints post-reactivation (**Figure 1c**). These confirmed our screen results, but we did note that some of the effects on ORF68 levels were not sustained at later timepoints, most notably when targeting TSSalt. This may reflect changing regulatory events as the viral lifecycle progresses. While guides targeting the EF1a promoter displayed the expected effect as well, these guides were highly toxic, most likely through their association with the BAC-encoded EF1a-EGFP-HygroR selection locus or the possible inhibition of the host EF1a site. Thus, we excluded this peak from further analysis.

Despite the correspondence of most peaks to TSSs, the large observed footprint of CRISPRi prevented high resolution identification of important underlying regulatory features. Illustrating this, the TSS68 peak is comprised of guides targeting not only the entirety of the ORF68 coding region, but also many surrounding genes (**Figure 1d**). This equates to approximately a 2-5 kb window of CRISPRi repression from a single guide, which is mirrored at other loci (**Figure S1d**). This large footprint is likely due to the spread of KRAB-induced heterochromatinization^37^ and should be taken into account when using sodium butyrate and CRISPRi on viral genomes, which display significantly higher gene density than cellular genomes. This prompted us to further interrogate the global and local transcriptional effects of CRISPRi at each regulatory region on the viral genome.

### CRISPRi recruitment to the viral genome inhibits many genes locally

The regulatory regions identified above could either impact ORF68 transcription specifically or could more globally disrupt KSHV lytic gene expression. To evaluate these possibilities, we performed polyA+ RNA-seq at 24 hours post-reactivation on the previously validated guide pools (**Figure S2a**). Silencing of the regulatory loci by CRISPRi caused changes in ORF68 mRNA levels that are consistent with those observed at the protein level by both RNA-seq and follow-up RT-qPCR (**Figure 2a, Figure S2b,c, Supplementary Data 3**), suggesting that regulation primarily controls RNA abundance. Nearly all viral genes were downregulated when ORF50, TSS75, or TSS57 were silenced, aligning with the critical roles of ORF50, ORF75, and ORF57 proteins in the viral lifecycle. ORF50 is a master regulator of KSHV lytic reactivation,^38^ ORF75 is required to prevent innate immune suppression of viral gene expression,^39,40^ and ORF57 has many reported viral functions including the export of viral mRNA from the nucleus.^41^

**Figure 2.**
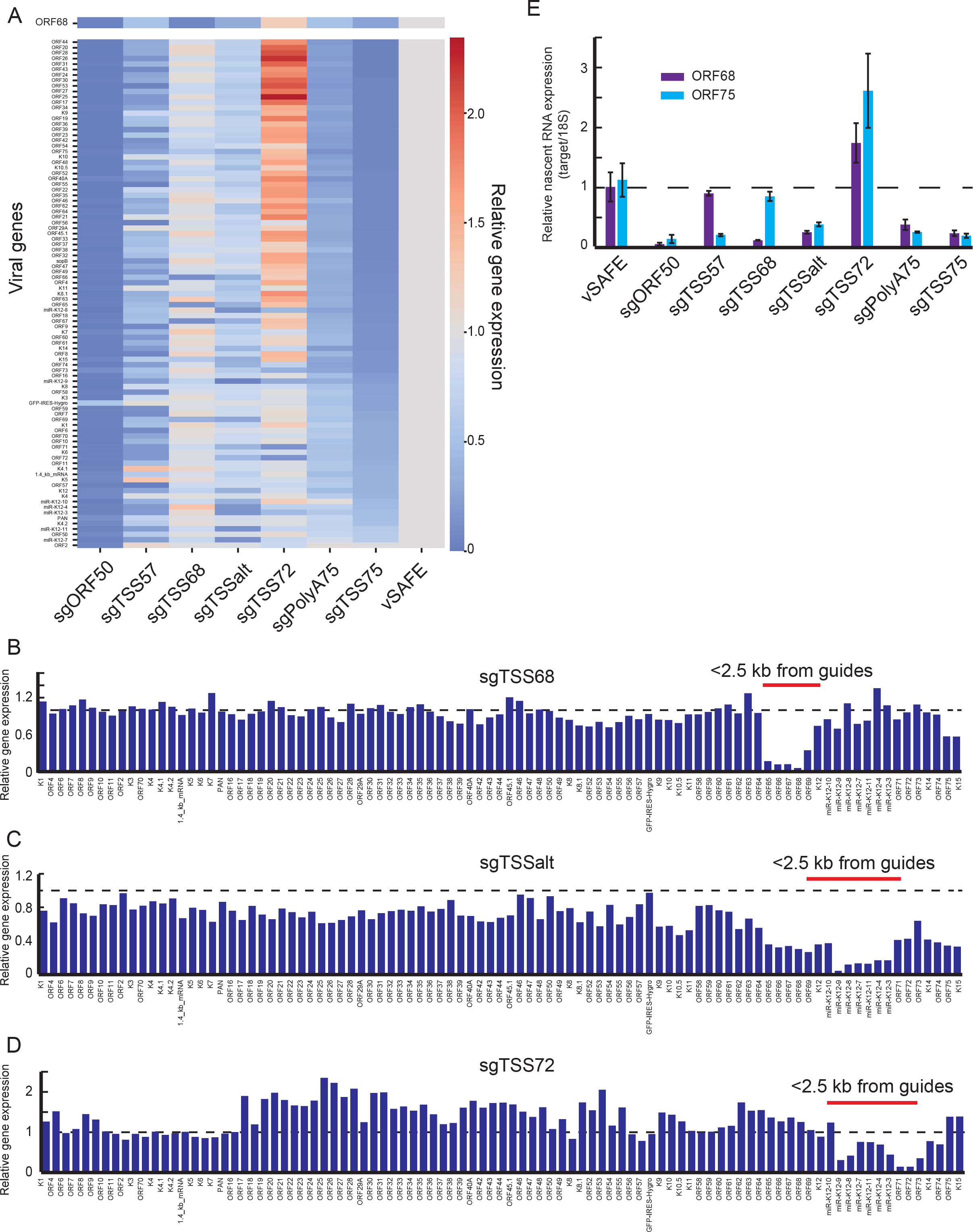
*Local effects caused by CRISPRi*. A) Heat map of changes to viral gene expression at 24 hours relative to vSAFE negative controls, with ORF68 presented at the top. Each column shows relative gene expression changes following CRISPRi-induced suppression of the indicated locus. Values are sorted by the effect of sgTSS75. Average of three replicates. B-D) Change in RNA-level of each viral gene relative to vSAFE negative controls in genome-order. Genes whose start codons are within 2.5 kb of at least one guide in targeting pool are highlighted. E) Nascent RNA expression was measured by RT-qPCR at 24 hours post-reactivation from cells treated for two hours with EU. Values are presented relative to parental cells, and error bars are standard error from three independent replicates.

In contrast, guides targeting TSS68 and TSSalt had a limited global effect, strongly inhibiting only a small number of genes each. The strongest downregulated genes fall within the local region of the guides (**Figure 2b,c**), reaffirming that CRISPRi inhibits transcription in region of approximately ∼2.5 kb around the targeting site. This appears true even when the effect of CRISPRi increases ORF68 mRNA expression, as in the case of TSS72. Silencing the TSS72 region causes a global increase in viral gene expression (**Figure 2a**) despite still locally decreasing transcription (**Figure 2d**). Many of these locally repressed regions include ORF73, which encodes for LANA, whose expression is required for latency maintenance,^3,42^ though whose knockdown was not seen here to affect ORF68 expression. To test whether targeting TSS72 is affecting latency, we grew our knockdown lines in the absence of hygromycin selection and observed a more rapid loss of latency only when targeting TSS75 or polyA75 (**Figure S2d**). Thus, RNA-seq corroborates our screening data, and reveals both local and global effects of CRISPRi at our target loci.

We next tested whether these altered ORF68 and ORF75 RNA levels stem from transcriptional changes after CRISPRi targeting of the regulatory loci. At 24 hours post reactivation, we used ethynyl uridine (EU) to metabolically label newly transcribed RNA for two hours. EU-labeled RNA was purified, modified with biotin, and isolated, and the levels of newly synthesized ORF68 and ORF75 mRNA were measured by RT-qPCR (**Figure 2e**). As expected, targeting TSS68 or TSS75 had a dramatic effect on their respective nascent RNAs, and silencing of other regulatory loci yielded a reduction or increase in EU-labeled RNA consistent with total RNA levels. The exception is TSS57, whose silencing did not significantly reduce nascent ORF68 mRNA at 24 hours post reactivation. Thus, while these regulatory loci all impact the total RNA abundance of their targets, they may do so through a combination of transcriptional and post-transcriptional mechanisms.

### Viral knockouts identify associated coding features

While CRISPRi at a given locus may effectively suppress multiple viral genes, not every gene will be responsible for the observed effect on ORF68 expression. For example, while guides targeting the TSS68 repress expression of ORFs 65, 66, 67, 68, and 69 (**Figure 2b**), downregulation of the ORF68 reporter is most likely due to direct repression of the ORF68 promoter. Furthermore, CRISPRi alone is unable to distinguish whether regulatory regions influence transcription of a regulatory protein or act by a non-coding mechanism. Therefore, we next directly assessed the role of coding loci underlying each regulatory locus by performing a CRISPR nuclease tiling screen, using a version of the HaloTag-ORF68 reporter iSLK line containing Cas9 instead of CRISPRi.

In previous screens, we have reported a strong background effect, where targeting any locus on the viral genome with Cas9 nuclease resulted in a decrease in reporter expression.^34^ Here, we observed unexpectedly that this background effect was variable across the viral genome, with targeting to the region upstream of the ORF68 coding region having a stronger effect on reporter expression than targeting downstream (**Figure S3a, Supplementary Data 4,5**). The reason for this difference is unknown (possibly a local effect of DNA damage on the viral genome), but to adjust for the differences in local background we used a boundary method to identify coding regions: for each candidate coding exon, we compared the enrichment of the coding region to the immediate adjacent non-coding region, and considered the coding region a hit if the boundaries were both significantly increased or both significantly decreased.

This yielded an exhaustive list of viral coding regions that control ORF68 expression at 24 hours post-reactivation: ORF50, HygroR, ORF68, and ORF75 (**Figure 3a-c; Figure S3b**). Targeting the corresponding CRISPRi loci of ORF50, TSS68, and TSS75 each disrupt ORF68 expression, while CRISPRi repression of the EF1a promoter controlling HygroR expression increased ORF68 expression. Loss of functional protein thus likely explains these regions’ effect on ORF68. In contrast, we observe no effect on ORF68 of targeting ORF57 and ORF72 coding regions with Cas9 (**Figure S3c,d**). Thus, TSS57 and TSS72 presumably impact ORF68 independently of their associated coding ORFs. The final two CRISPRi loci, TSSalt and PolyA75, are not located at the promoters of any coding regions, and indeed we find no evidence of a coding region near TSSalt that affected ORF68 expression. PolyA75 is adjacent to the ORF75 coding locus, but no other coding elements are identified that could explain its activity. These regions with no associated coding region thus likely act through noncoding elements that are unable to be efficiently disrupted by Cas9.

**Figure 3.**
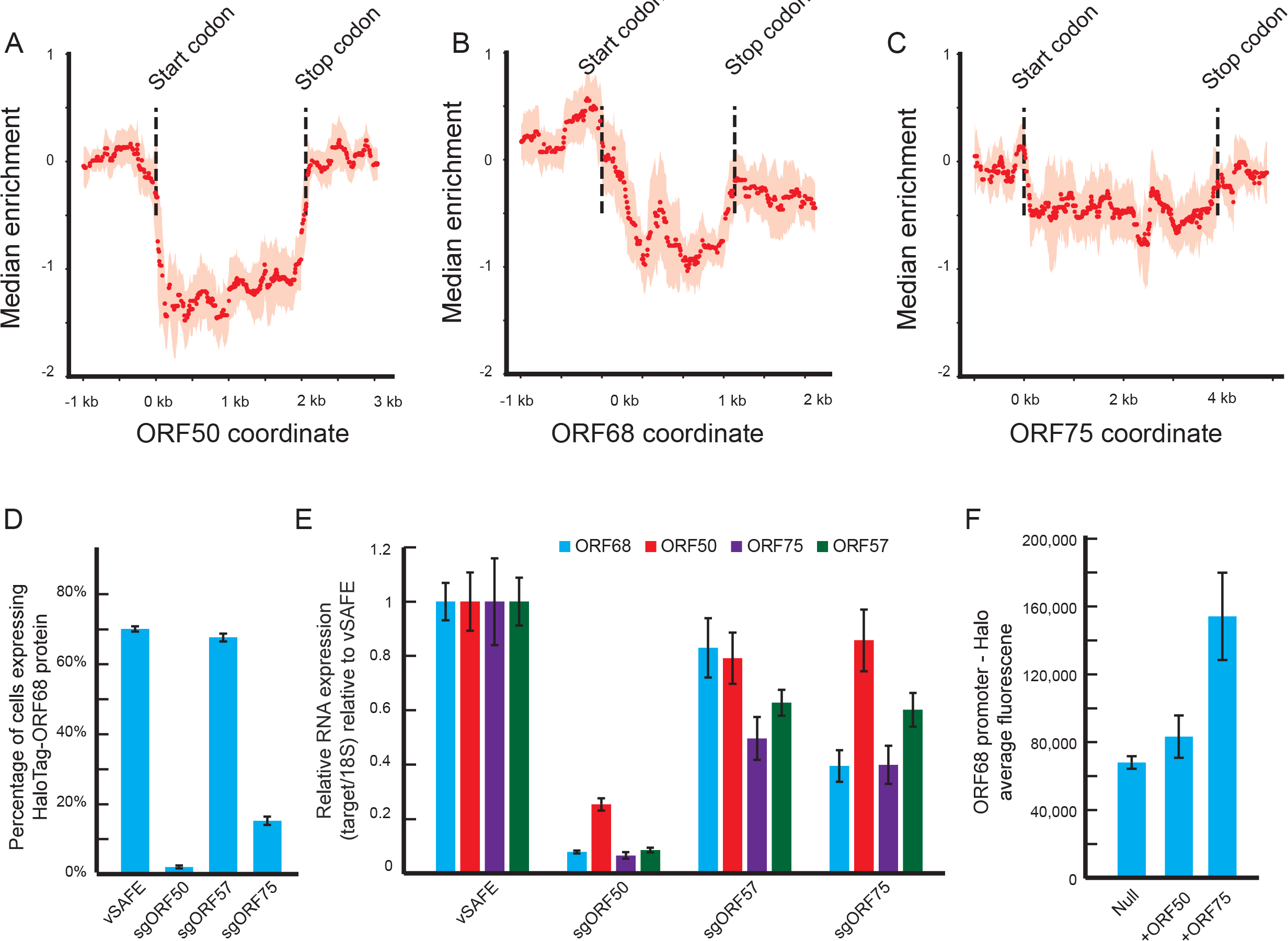
*Knockout screen maps associated coding regions*. A-C) Median smoothed enrichments from Cas9 nuclease screen at associated coding locus. Dotted lines indicate exon boundaries. For each guide, the median enrichment of 500 bp window centered at the target locus was calculated along with an IQR. Median value is shown as a point and IQR is shown as shaded region. Regions were considered significant if the guides on each side of the boundary were significantly different and in consistent directions. D) Percent of cells expressing HaloTag-ORF68 24 hours post-reactivation for the indicated pool of coding region-targeting guides. Values are averages of four independent replicates and error bars represent standard error. E) RT-qPCR measurements of ORF68, ORF50, ORF75 and ORF57 mRNA at 24 hours post-reactivation following Cas9-based targeting of the loci indicated on the x-axis. Data are presented relative to 18S RNA and vSAFE. Error bars show standard error from four technical replicates. F) Average fluorescence from HEK293T cells transfected with an ORF68-promoter-driven HaloTag and a plasmid expressing the indicated viral protein. Error bars are standard error from seven independent replicates.

To test whether the identified coding regions have the expected negative effect on the mRNA levels of ORF68, we cloned pooled guides for nuclease-targeting at the coding regions of ORF50 and ORF75, including ORF57 as a negative control. ORF50 and ORF75 pools had the expected effect of decreasing ORF68 protein levels (**Figure 3d**) as well as corresponding depletion of ORF68 RNA (**Figure 3e**). To determine whether ORF50 and ORF75 are direct or indirect activators of ORF68, we transfected a reporter plasmid containing 230 bp of upstream of the ORF68 start codon driving a HaloTag reporter along with an expression plasmid of either ORF50 or ORF75 into HEK293T cells. Only ORF75 caused a significant increase in expression from the ORF68 promoter (**Figure 3f**). Given that ORF50 is required for ORF75 expression (**Figure 3e**), these data suggest a regulatory model of protein coding elements where ORF50 activates ORF75 which, in turn, activates ORF68. We also observed that targeting either ORF50 or ORF75 reduced RNA levels of ORF57 (**Figure 3e**). Interestingly, sgORF57 pools reciprocally decreased ORF75 RNA levels, yet ORF57 protein disruption had little or no effect on ORF68 mRNA levels – as predicted by the nuclease screen – suggesting an unclear regulatory relationship.

### Mapping regulatory targets of noncoding elements

The remaining CRISPRi peaks at TSS57, TSSalt, TSS72, and polyA75 presumably are noncoding loci that instead act in a distal manner to indirectly impact one of the three viral proteins that affect ORF68 expression. We therefore returned to our RNA-seq data and measured the correlation between each sample (**Figure 4a**). We hypothesized that despite local effects of CRISPRi, the regulatory regions would correlate most strongly with their regulatory target, and as we have identified all regulatory regions the number of potential targets is limited. TSSalt and polyA75 most strongly correlated with TSS68 and TSS75 respectively, suggesting that these regions promote ORF68 and ORF75 expression. Conversely, TSS72 had a strong anticorrelation with polyA75 and TSS75. Given that recruitment of CRISPRi to TSS72 causes an increase in viral gene expression, this suggests that TSS72 acts to repress expression of ORF75. TSS57 weakly correlated with many targets, preventing any firm conclusion. We can thus use these correlative regulatory interactions along with data from **Figure 3** to create a model regulatory network consisting of both coding and non-coding elements controlling ORF68 expression (**Figure 4b**). Of note, as we performed these experiments while overexpressing ORF50 from an exogenous promoter, we were likely unable to detect any regulation of the viral copy of ORF50. While silencing TSS57, ORF57, or TSSalt each decreased transcription of ORF75 (**Figure 2e,f**; **Figure 3e**), we have excluded these interactions from our model as we did not observe the expected subsequent changes to global or ORF68 transcription (**Figure 2a**).

**Figure 4.**
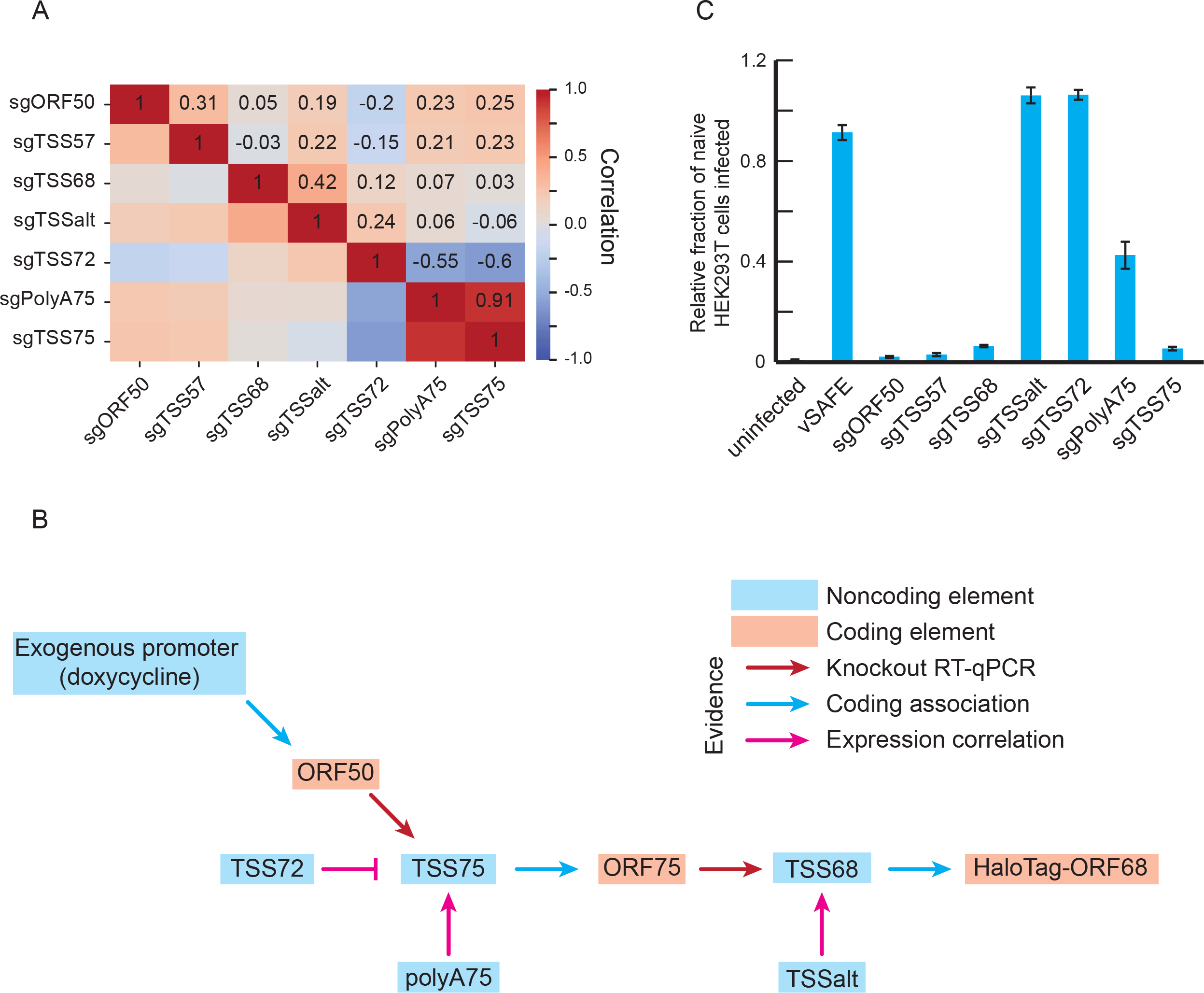
*Mapping regulatory network by effect on viral transcription*. A) Co-correlation matrix of the RNA-seq data from Figure 2a. For each indicated pair of sgRNA pools, a Pearson correlation was calculated between the viral RNA levels. B) Model of regulatory events controlling transcription of the ORF68 locus. C) Supernatant transfer assay measuring changes in KSHV virion production after knockdown of the indicated loci. Error bars represent standard error from six independent reactivations.

To test this model, we evaluated how the components of this ORF68 regulatory network impacted virion production in KSHV infected cells using a supernatant transfer assay. BAC16 derived KSHV expresses GFP, which enables quantification of infected recipient HEK293T cells by flow cytometry. As expected, targeting any of the coding elements via their promoters (TSS57, TSS68, TSS75) caused a severe loss in infectious virion production (**Figure 4c**). Targeting polyA75 also reduced virion production, albeit more modestly, consistent with the regulatory network. In contrast, guides targeting TSS72 or TSSalt did not negatively impact virion production (**Figure 4c**). This was expected for TSS72, whose silencing increases ORF68 expression. However, that targeting TSSalt – which specifically disrupts ORF68 expression at the early 24h but not the late 48h late time point (see **Figure 1c**) – did not impair virion production suggests that while ORF68 expression is essential late in infection, it may be dispensable at early time points.

### Physical interactions of noncoding regions and their targets

We hypothesized that these noncoding elements at TSSalt, TSS72, and polyA75 function distally as either enhancers or regulators of the 3D viral genome structure. Enhancers are expected to be physically close to their target promoters, which can be determined by measuring genome architecture using chromosome capture techniques like Hi-C. Previous work has demonstrated that high-quality viral-viral or viral-host contacts can be obtained using capture Hi-C.^5,43,44^ We thus evaluated this approach on iSLK-BAC16 cells 24 hours post reactivation (**Figure 5a; Supplementary Data 7**). Indeed, by applying KSHV-specific sequence capture to our libraries, we were able achieve high (<2 kb) resolution contact frequency maps on the viral genome (**Figure S5a**) with relatively few sequencing reads (∼7 million reads). Contact features were clear at 1-2 kb resolution but still notable at even higher (500 bp) resolution (**Figure S5a**).

**Figure 5.**
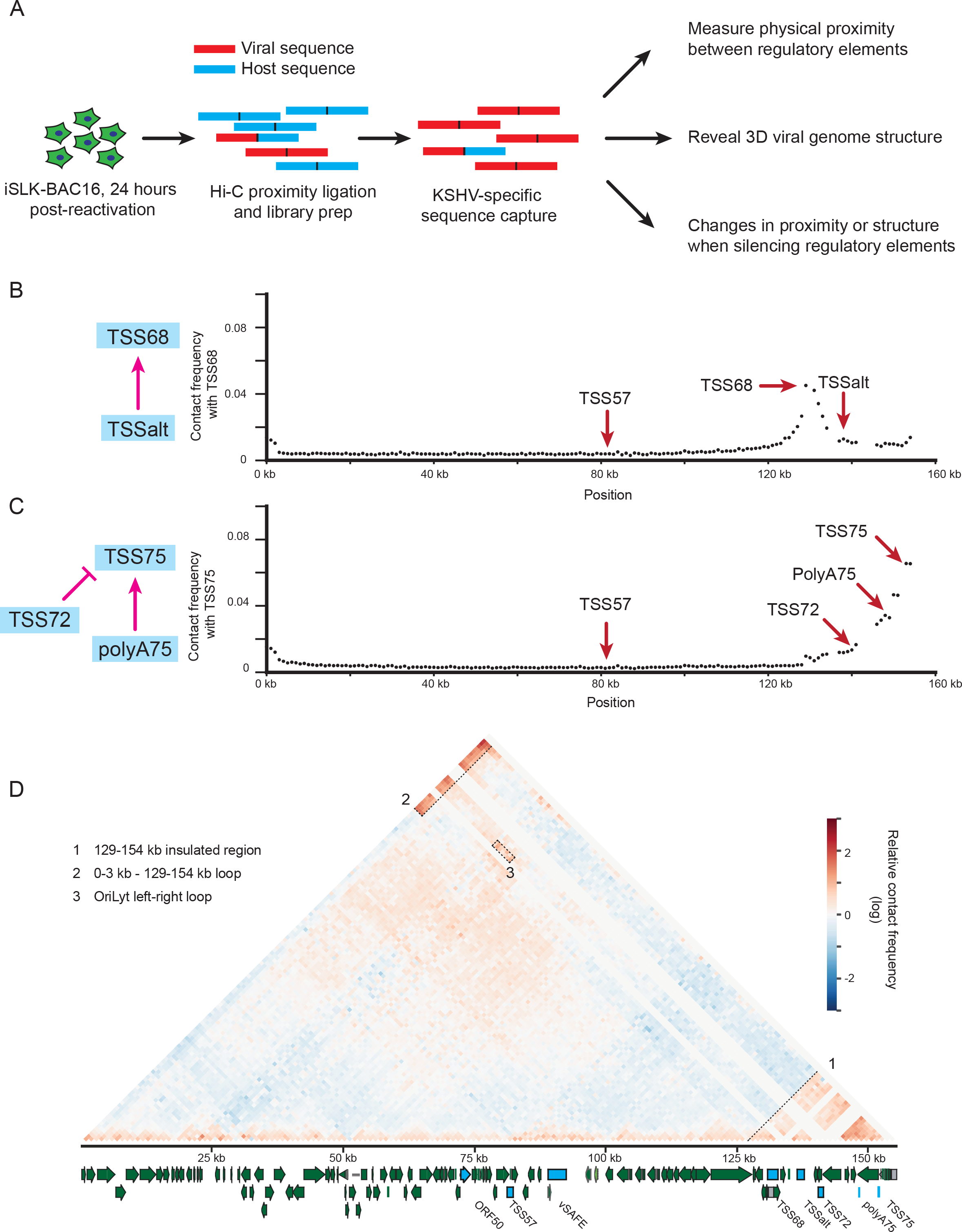
*Capture Hi-C of the KSHV genome*. A) Schematic of capture Hi-C experiment. B,C) Contact frequency between B) TSS68 and C) TSS75 and other locations in the viral genome at 1 kb resolution. D) Relative contact frequency map corrected for circular/concatenated distance. Noted features are marked and labeled. Positive values represent more interaction than expected. Annotated viral genome is provided with regulatory elements identified marked in blue.

We first asked whether the regulatory elements that interact functionally are also physically proximal, which would be consistent with their activity as enhancers. To function as an enhancer, TSSalt should have a high contact frequency with its target TSS68, while TSS72 and polyA75 should have a high contact frequency with their target TSS75. Indeed, this is the case: when looking at the contact frequency of these regions with the rest of the viral genome at 1 kb resolution, TSSalt, TSS72 and polyA75 are physically close to their identified targets but not to another regulator region, TSS57 (**Figure 5b,5c**). This also holds true in the reciprocal interactions (**Figure S5b-d**). These results are consistent with TSSalt, TSS72, and polyA75 functioning as enhancers and exclude the possibility that TSS57 itself is acting as an enhancer targeting either TSS68 or TSS75.

We next asked whether this physical proximity is greater than would be expected given the linear genomic distance between the elements and their regulatory targets. Previous Hi-C data has consistently identified that contact frequency is reduced with distance and that this decay can be modeled using a power law.^45,46^ However, the relationship between linear distance and the observed contact frequency on the viral genome diverged at higher distances (**Figure S5e**), presumably reflecting that the KSHV genome is present in circular and concatenated forms.^43^ Indeed, a distance metric adjusted for these forms better matched the power law expectation (**Figure S5f**) and was used for subsequent analyses.

By normalizing the measured contact frequencies (**Figure S5a**) to the expected contact frequency based on distance, we obtained a relative contact frequency that represents the structure of the viral genome at 1 kb resolution (**Figure 5d**). In agreement with the close contacts shown in **Figure 5b-c**, we observed a strongly associated region from ∼129-154 kb that contains TSS68, TSSalt, TSS72, polyA75, and TSS75 (**Figure 5d, outlined region 1**), indicating that these regulatory elements are physically closer than expected given their linear distance. Of note, this corresponds to the region with increased background in the Cas9 screen (**Figure S3a**), consistent with disruptions to this region causing changes to local transcription. Compartments like this exist on the KSHV genome at a smaller scale than typical transcriptional-associated domains (TADs),^43^ so we instead refer to them as insulated regions. Other stand-out physical associations include the 129-154 kb insulated region and the 1-3 kb region containing promoters for K1, ORF4, and ORF6 (**Figure 5d, region 2**), as well as the region around TSSalt (which includes the right lytic origin of replication), and region at ∼24 kb which includes the left lytic origin of replication (**Figure 5d, region 3**). These physical associations may represent chromosomal loops between distal regions of the viral genome. However, evaluating whether these loops are statistically significant or if there are additional loops present would require a robust statistic, and it is unclear what would be appropriate given the small size of the viral genome, the high frequency of distal interactions, and the irregular shape of the loops we do see.

### Changing physical relationship between regulatory regions

If ongoing transcription at these regulatory elements is important for physical interaction with their targets, we reasoned that silencing them could alter the architecture of the viral genome. We tested this by targeting each regulatory element using CRISPRi, then performing capture Hi-C to measure how the 3D structure of the viral genome changed. We modeled how the genome structure changes by grouping the resulting capture Hi-C maps into initial, intermediate, and final stages, representing progression through the viral lytic lifecycle and the regulatory network controlling ORF68 expression by introducing a roadblock caused by silencing each element (**Figure 6a; Supplementary Data 7**). Maps from unreactivated cells and reactivated cells with silenced ORF50 (sgORF50) represented the initial stage, as the viral lytic cycle cannot progress in the absence of ORF50. The intermediate stage comprised the maps from reactivated cells with silenced TSS75, TSS57, and PolyA75, as the RNA-seq data indicated these cause significant changes to the viral transcriptome. The final stage comprised the maps from cells with silenced TSS68, TSSalt, the negative control vSAFE, and the negative regulatory element TSS72.

**Figure 6.**
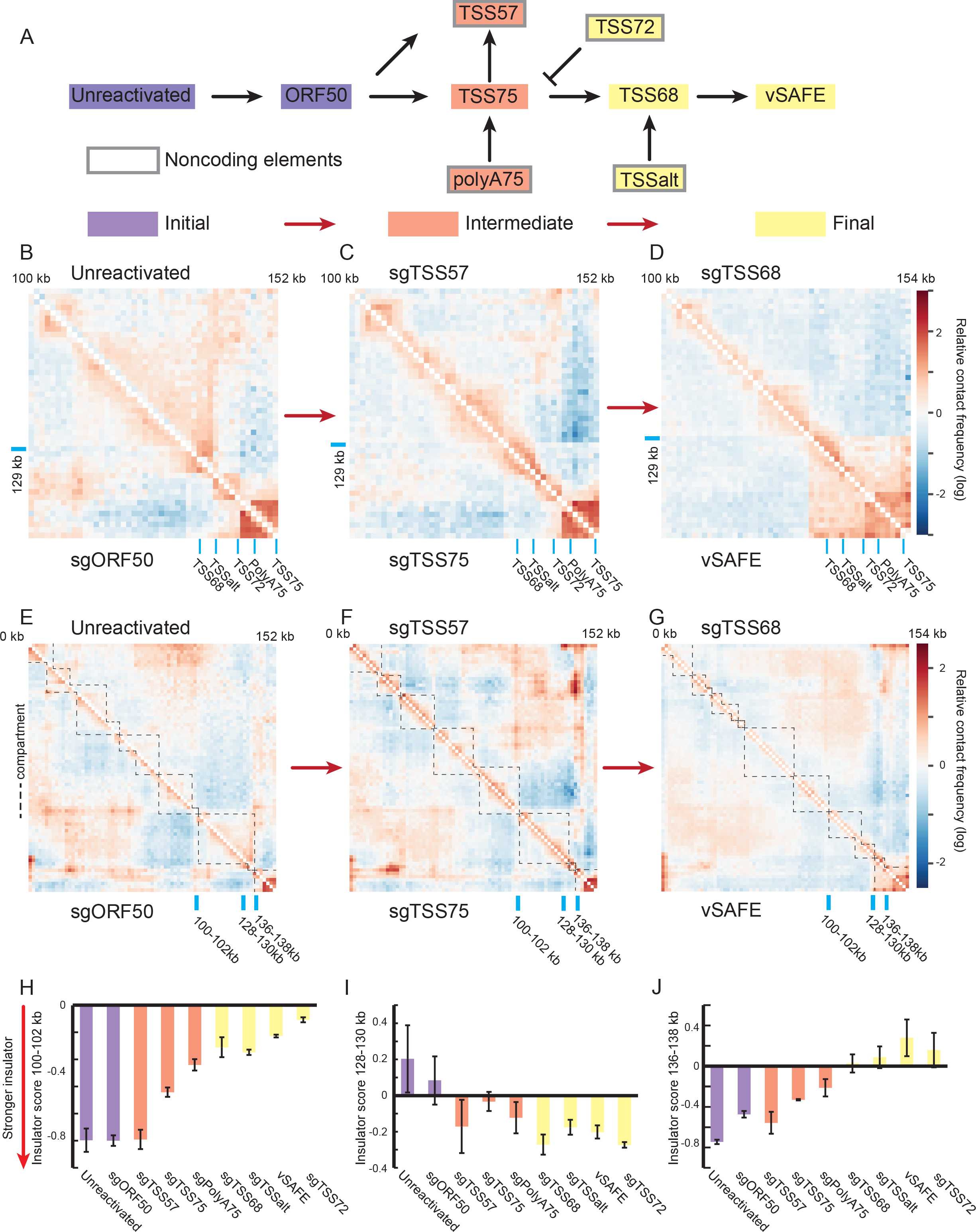
*Changing physical relationship between regulatory regions.* A) Schematic splitting the regulatory network into initial, intermediate, and final stages. B-D) Relative contact frequency at 1 kb resolution from 100-154 kb measured by capture Hi-C 24 hours post-reactivation for representative B) initial stages (purple), C) intermediate stages (orange), D) and final stages (yellow). Positive values indicate greater interaction than expected, and locations of regulatory elements are marked in blue. E-G) Relative contact frequency at 2 kb resolution across the genome measured by capture Hi-C 24 hours post-reactivation for representative E) initial stages, F) intermediate stages, G) and final stages. Dotted lines represent locations of insulator regions as defined by negative local-minimum insulator scores. The locations of three observed insulators are marked in blue. H-J) Insulator scores at the marked locations in A-C at D) 100-102 kb, E) 130- 132 kb, and F) 136-138 kb. More negative values indicated a stronger insulator. Values are the average of two adjacent regions with error bars representing standard deviation of these two values.

We first examined the insulated region at 129-154 kb, which contains many of these regulatory elements. At 1 kb resolution, we observed large structural changes to this region as the viral life cycle progresses from the initial (**Figure 6b**) to the intermediate (**Figure 6c**) to the final stage (**Figure 6d**), with this 129-154 kb compartment only clearly forming in the final stage. We quantitatively assessed these structures using an insulator score, which was calculated from the relative frequency of contacts across a given region. Insulators “protect” expressing genes from the neighboring genomic environment; they were thus detected here as areas with fewer than expected contacts that span the region, i.e., a low insulator score. These values showed strong concordance within each stage at 2 kb resolution (**Figure S6a-c**), and we used them to visualize compartments by defining their surrounding insulator boundaries as regions with locally minimum insulator scores across the viral genome (**Figure 6e,f,g)**. These results are consistent with previous work demonstrating that the KSHV genome is highly structured before and after reactivation.^43,47–49^ However, we note that the insulator strength in the final stage is globally reduced (**Figure S6c**), supporting reduced chromatinization of the viral genome in later stages of the lytic cycle.^2^ We did not observe any large shifts in 3D structures when silencing polyA75, TSSalt, or TSS72 when compared to other targets in their stages (**Figure S6d-f**), suggesting they only regulate the 3D genome structure via their regulatory targets.

In addition to observing changes in the compartment structure between stages, we also quantified the strength of the insulators at a given region that separates these compartments. As the regulatory circuit progresses, there is a weakening of the insulator at 100-102 kb, an appearance of an insulator at 128-130 kb, and the loss of an insulator at 136-138 kb (**Figure 6h-j**). This corresponds to the region at 128-138 kb moving from a compartment at 100-130 kb to form the ∼129-154 kb insulated region we originally noted (**Figure 5d**). As this region contains ORF68, we next asked how these changes affect the contact frequency between the regulatory elements and their targets. While we do not see any large shifts in the physical interactions of the ORF72 locus with TSS75 relative to the reactivated control (**Figure 6g**), the interaction between polyA75 and TSS75 is stronger in both the initial and intermediate stages (**Figure 6h**). This is consistent with our functional data showing that targeting polyA75 reduces ORF75 function even before reactivation (**Figure S2d**) and may represent a dilution of the interactions as the insulator at 136- 138 kb is lost (**Figure 6j**) and the compartment containing polyA75 and TSS75 is expanded (**Figure 6f,g**). In contrast, we find that TSSalt and TSS68 are farther apart when targeting either TSS75 or TSS57 (**Figure S6i**), suggesting that this regulatory interaction is inhibited when silencing these regions. While TSS75 likely also acts directly on TSS68 via the ORF75 protein (**Figure 3f**), this allows us to complete the regulatory network by incorporating the effect of targeting TSS57 on the interaction between TSS68 and TSSalt (**Figure S6j**). Thus, by silencing individual regulatory elements and performing capture Hi-C, we determined how each element contributes to the 3D structure of the viral genome; by further incorporating the model for ORF68 transcriptional regulation, we also mapped the 3D changes to the genome as the viral lifecycle progressed.

## Discussion

Here, we identified novel regulatory events across the KSHV genome that control the expression of ORF68 and were able distinguish between coding and noncoding elements, identify their regulatory targets, and establish their changing physical relationships in the 3D genome. Like many other viral early genes, ORF68 has a simple expression pattern: it turns on early in the lytic cycle and remains on throughout. In this light, the model regulatory network we built – and characterized through exhaustive functional genomics, transcriptional profiling, and capture Hi-C – is surprisingly complex. The virus borrows widely from the regulatory toolbox of its human host, spanning promoters, enhancers, repressors, and 3D structural elements. However, unlike the host, all these elements are contained within a compact, densely encoded genome, allowing us to identify a near-complete architecture controlling gene expression within the human nucleus.

Our functional model does not make predictions for the mechanistic nature of all these regulatory events. For example, the coding elements may or may not directly regulate transcription on the viral genome. Indeed, whereas ORF50 has been previously characterized as a viral transcription factor,^38,50^ ORF75 more likely acts indirectly by preventing inhibition of viral transcription by host factors.^51^ Similarly, the non-coding elements could involve ncRNAs, microRNAs, or enhancers, as CRISPRi is expected to be able to inhibit each of these; though as we see a direct effect on transcription (**Figure 2b)** and physical proximity between regulatory region and target (**Figure 5**), we favor an enhancer model. Given the density of the viral genome and the width of CRISPRi footprint, it is possible a peak corresponds to multiple regulatory events or that two nearby peaks both inhibit the same regulatory locus. The TSSalt here could be an example of the former, as there are nearby K12 TSS and miRNA loci (**Figure S1e**), and indeed a recent investigation has proposed that two distinct sequences here have regulatory activity.^52^ The polyA75 peak could be an example of the latter as it behaves very similarly to the nearby TSS75 peak. Though with the viral density and propensity for multiple functions encoded within the same locus, these issues may be intrinsic to the study of gene regulation on the viral genome, but it is possible that other functional perturbations such as using base editors or dCas9 could allow us to pinpoint the underlying regulatory sequence.

It is also difficult to distinguish the effects of disrupting ORF57 protein and the regulatory element at TSS57. Our observation that Cas9-based disruption of ORF57 did not affect ORF68 mRNA or protein suggests that ORF57 protein does not regulate ORF68. This is in agreement with previous reports using an ORF57 deletion virus, which also indicated little regulatory effect on ORF68.^53^ ORF57 protein does have a broad effect on viral transcription, but this is likely through direct effects on DNA replication components, upon which the majority (but not ORF68^30^) of gene expression is partially dependent. It is possible that the TSS57 promoter contributes to ORF68 regulation independent from the ORF57 protein, as many human promoters act as enhancers for distal genes,^54^ but we do not observe a physical association between TSS57 and either TSS75 or TSS68 (**Figure 5b,c**). Instead, we see that targeting TSS57 causes a large disruption to the 3D conformation of the KSHV genome (**Figure 6c**), leading to a reduced interaction between TSSalt and TSS68 (**Figure S6i**), potentially disrupting this enhancer-promoter relationship. However, this remains a hypothesis as we cannot disrupt the promoter and its hypothetical structural activity using CRISPRi without also disrupting the protein levels of ORF57.

Our approach demonstrates the power of combining CRISPR screening tools for the discovery of viral gene regulatory networks. Given that KSHV is a nuclear double-stranded DNA virus, it incorporates the same spectrum of regulatory mechanisms as the host – transcription factors, non-coding RNAs, enhancers, and DNA structural elements. By studying these networks on the viral genome, we can thus learn both how these regulatory mechanisms function and, ultimately, how the virus controls gene expression under diverse cellular conditions and in various cell types.

## Methods

### Plasmid and oligos

pMD2.G (Addgene plasmid # 12259), pMDLg/pRRE (Addgene plasmid # 12251) and pRSV-Rev (Addgene plasmid # 12253) were gifts from Didier Trono. pMCB320 was a gift from Michael Bassik (Addgene plasmid # 89359). lentiCas9-Blast was a gift from Feng Zhang (Addgene plasmid # 52962). Lenti-dCas9-KRAB-blast was a gift from Gary Hon (Addgene plasmid # 89567). Sequences used are listed in **Supplementary Data 6**.

### Cell culture and plasmids

iSLK^55^ and Lenti-X 293T (Takara) cells were maintained in DMEM (Gibco +glutamine, +glucose, -pyruvate) with pen-strep (Gibco; 100 I.U./mL) and 1X Glutamax (Gibco) along with 10% FBS (Peak Serum). iSLK cells were maintained in 1 ug/mL puromycin, 50 ug/mL G418, and 125 ug/mL hygromycin B (Gibco). Cas9+ and CRISPRi+ cells were additionally maintained in 10 ug/mL blasticidin. 0.05% Trypsin (Gibco) was used to passage cells. All cells were maintained at 37C and 5% CO2 in a humidity-controlled incubator. Lenti-X 293T cells were obtained from the UCB Cell Culture Facility.

### Generation of CRISPRi Halo-ORF68 iSLK line

iSLK cells latently infected with a copy of BAC16^56^ containing a HaloTag-ORF68 fusion at the endogenous locus were lentivirally infected with dCas9-KRAB (CRISPRi). Briefly, Lenti-X cells were transfected with third-generation lentiviral mix (pMDLg/pRRE, pRSV-REV, pMD2.G) and dCas9-KRAB (blastR) with polyethylimine (Polysciences). Supernatant was harvested at 48 and 72 hours and 0.45 um filtered before applying to iSLK cells for 48 hours. This process was then repeated to ensure high CRISPRi expression.

### CRISPRi screening and analysis

A library of guides tiling the KSHV BAC16 genome was delivered lentivirally to the CRISPRi+ iSLK cells above. Four days later cells were reactivated with 5 ug/mL doxycycline and 1 mM sodium butyrate and treated with 10 nM JF646 Haloligand (Promega). 24 hours post reactivation, cells were fixed in 4% PFA, and sorted for high and low ORF68 expression using a BD Aria II. Cells were then unfixed overnight in 150 mM NaCl and 60C and 50 ug/mL protease K (Promega). DNA was then extracted using a single column of QIAamp DNA Blood Mini Kit (Qiagen), following the manufacturers’ protocol and adjusting initial reaction volume. The sgRNA locus was then amplified and library adaptors ligated as previously described^57^. Libraries were sequenced on an Illumina NovaSeq 6000.

Counts for individual guides were converted to enrichment scores by calculating the log2 ratio of counts between high and low populations relative to the median negative control. Enrichment values from two replicates were averaged. To calculate significance of a given window, a 500 bp sliding window was used, comparing the enrichment of each guide and a 500 bp neighborhood to the enrichment scores of all negative controls using a Mann-Whitney test. An arbitrary p-value cutoff was used to identify peaks.

### Individual guide delivery

For each pool of sgRNAs, independent lentiviruses were produced as above, then pooled and applied to CRISPRi-positive HaloTag-ORF68 iSLK cells for 48 hours. For protein analysis, cells were reactivated in the presence of 10 nM JF646 Haloligand (Promega) with doxycycline and sodium butyrate as above. Cells were then analyzed for fluorescence at 24 and 48 hours post- reactivation from four independent reactivations on a BD Accuri C6 plus.

For latency analysis, cell lines were maintained in triplicate with blasticidin, puromycin, and G418 but in the absence of hygromycin. Cells were split every 48 hours and GFP levels were measured by a BD Accuri C6 plus. The percent of GFP positive cells on day 11 was normalized to the percent positive on day 1, and this ratio was again normalized to the loss of GFP observed in parental cells without a guide RNA.

### RNA-seq and analysis

RNA samples from above were sent for library preparation and sequencing at the QB3-Berkeley Genomics core labs (RRID:SCR_022170). Total RNA quality as well as poly-dT enriched mRNA quality were assessed on an Agilent 2100 Bioanalyzer. Libraries were prepared using the KAPA RNA Hyper Prep kit (Roche KK8581). Truncated universal stub adapters were ligated to cDNA fragments, which were then extended via PCR using unique dual indexing primers into full length Illumina adapters. Library quality was checked on an AATI (now Agilent) Fragment Analyzer. Library molarity was measured via quantitative PCR with the KAPA Library Quantification Kit (Roche KK4824) on a BioRad CFX Connect thermal cycler. Libraries were then pooled by molarity and sequenced on an Illumina NovaSeq 6000 S4 flowcell for 2 x 150 cycles, targeting at least 25M reads per sample. Fastq files were generated and demultiplexed using Illumina bcl_convert and default settings, on a server running CentOS Linux 7.

Sequencing quality was assessed with MultiQC and reads were preprocessed with HTStream version 1.3.0 including deduplication. Genome indices were prepared using STAR 2.7.1a. The human GRCh38.p13 genome assembly was indexed with Gencode v43 annotations. Due to overlapping transcripts on the KSHV-BAC16 genome, individual exon coordinates were assigned to the corresponding parent transcript. Preprocessed reads were then aligned using STAR and counts files were generated for transcripts. Any transcript with no reads in all replicates and conditions was eliminated from further analysis. Reads from *E.coli* genes on the BAC were also removed. Raw viral counts were normalized to total input reads for each sample and subsequently normalized to within replicate vSafe condition values. Correlations and heatmaps were generated using the matplotlib, pandas, and seaborn packages in Spyder 5.3.3.

### RT-qPCR and EU-RT-qPCR

For RT-qPCR analysis, RNA was extracted at 24 hours post-reactivation using a RNeasy Plus Micro kit (Qiagen), treated with DNAase I (Lucigen), and reverse transcribed using AMV RT (Promega) and random 9-mers in the presence of RNasin (Promega). qPCR was then performed on a Quantstudio 3 using the indicated targets and with iTaq Universal SYBR Green (BioRad). RQ values were calculated using a standard curve. Results are from four independent reactivations.

For nascent RNA measurements, cells were reactivated, and 24 hours post-reactivation were treated with 200 uM 5-ethynyl uridine (EU) for two hours. RNA was then extracted using the RNeasy Micro Plus Kit (Qiagen) and EU incorporated RNA was labeled and purified per the Click- iT™ Nascent RNA Capture Kit (Invitrogen). Reverse-transcription was performed on-bead using SuperScript™ VILO™ cDNA Synthesis Kit (Thermo) reverse transcriptase, and qPCR was performed with iTaq Universal SYBR Green (BioRad) or PowerUp™ SYBR™ Green Master Mix (Thermo) on a Quantstudio 3 (Thermo). Relative quantities were calculated using a standard curve. Results are from three independent reactivations.

### CRISPR nuclease screen and analysis

Cas9 screen was performed as the CRISPRi screen above using a Cas9+ iSLK cell line infected with a copy of BAC16 containing a HaloTag-ORF68 fusion. The library was then amplified using staggered primers^58^ and a modified amplification protocol previously described.^59^ qPCR was used to determine cycle number at ¼ CT. All PCR product was run over a single Minelute column (Qiagen) for each sample. Size selection was performed using a gel extraction (Thermo). Library was sequenced on Illumina NextSeq 2000 with a 150bp single-read using Illumina sequencing primers. Adaptors were removed using cutadapt,^60^ and reads were aligned to library using bowtie.^61^

Log2 values were calculated as above, and enrichment values were averaged from two replicates. For each exon boundary, a window of 500 bp on one side was tested against a 500 bp on the other to calculate p-values using the Mann-Whitney test. Median values were used to determine the sign of the shift. Exons with p values < 0.001 and consistent signs were considered hits.

### Transfection of ORF68 reporter

HEK293T cells were transfected using PolyJet In Vitro DNA Transfection Reagent (SignaGen) with a plasmid containing 240 basepairs upstream of ORF68’s start codon^32^ driving a HaloTag. Cells were cotransfected with either a plasmid expressing ORF75 or ORF50. 24 hours later, cells were treated with JF646 Halo Ligand (Promega) for 24 hours before analysis on a BD Accuri C6 plus. Cells were then gated for JF646 signal, and the average fluorescence for the 646 positive cells was calculated. Data are from seven independent replicates.

### Supernatant transfer

Cells were reactivated with 5 ug/mL doxycycline and 1 mM sodium butyrate. 72-hour post reactivation, supernatant was filtered through a 0.45 um PES filter and applied to naïve HEK293T cells for 24 hours. HEK cells were then counted on a Accuri C6 Plus (BD) and percent GFP positive was used to calculate infection.

### BAC16-specific Capture Hi-C

Capture Hi-C data was generated using the Arima-HiC+ kit, a custom Arima panel specific to the BAC16 KSHV genome, and the Arima Library Prep Module according to the Arima Genomics manufacturer’s protocols. Briefly, 24 hours post reactivation, cells were fixed in 4% PFA and frozen. Chromatin was then purified, digested, and filled-in with biotin-labeled nucleotides. Proximity ligation was performed, DNA was purified and sheared, and ligated molecules were enriched by biotin pulldown. Library preparation was then performed, and viral-specific sequences were enriched using capture probes. Libraries were then sequenced using a NextSeq 2000 P2 150PE kit (Illumina).

### Capture Hi-C analysis

Ditags for Hi-C analysis were aligned to the BAC16 KSHV genome including the HaloTag insertion and filtered using HiCUP version 0.8.0^62^ and converted to fragment level counts using CHICACO bam2chicago.^63^ Fragments corresponding to the terminal repeats were removed before analysis. The viral genome was then split into bins of 0.5 – 5 kb, and each fragment was assigned to a bin based on the center of the fragment. Each bin was removed if it had less than a total of 5000 associated ditags. Bin level counts were then normalized to a symmetric stochastic matrix to calculate contact frequencies by iterative normalization of both rows and columns (Sinkhorn-Knopp). Distance to contact frequency relationship was then graphed using a linear distance metric, and a power law was fit. A second distance metric assuming a circular or concatenated genome was then used, allowing for a 180 kb genome size including ∼30 terminal repeats; if the linear distance was greater than half the genome size, then the genome size minus the linear distance was used. A power law was fit to this distribution, and relative contact frequencies are represented as the log ratio of the observed vs the expected power law distribution.

Insulator scores were calculated using the principles presented in Crane et al 2015.^64^ Contact frequencies spanning each region at 6-10 kb were averaged and normalized to the median value across the genome. Local minima of values lower than 0 were used as insulator locations.

## Supplementary Data

*Supplementary Data 1. CRISPRi count files.* Raw counts for sequencing from CRISPRi screens.

*Supplementary Data 2. CRISPRi screen results.* Processed values for CRISPRi screen.

*Supplementary Data 3. RNA-seq counts.* Viral counts for RNA-seq.

*Supplementary Data 4. Cas9 count files.* Raw counts for sequencing from Cas9 screens.

*Supplementary Data 5. Cas9 screen results.* Processed values for Cas9 screens.

*Supplementary Data 6. Sequences.* Sequences used.

*Supplementary Data 7. Hi-C results*. Binned interaction counts for each condition.

## Conflict of interest

The authors declare that they have no conflict of interest.

## Acknowledgments

The authors would like to thank the Xiaowen Mao and the members of the Glaunsinger lab for their feedback and support, and Dr. C. Kimberly Tsui for the stagger primer sequences. Cell lines were obtained from the UCB Cell Culture Facility which is supported by The University of California Berkeley (SCR_017924). RNA-seq was performed by the Berkeley functional genomics core (QB3 Genomics, UC Berkeley, Berkeley, CA, RRID:SCR_022170). Flow cytometry and FACS were conducted at the CRL Flow Cytometry Facility. We thank Hector Nolla and Alma Valeros of the UC Berkeley Cancer Research Laboratory Flow Cytometry Facility for training and expertise. This work used the Vincent J. Coates Genomics Sequencing Laboratory at UC Berkeley, supported by NIH S10 OD018174 Instrumentation Grant. B.A.G. is an investigator of the Howard Hughes Medical Institute, and D.W.M. is a Howard Hughes Medical Institute Awardee of the Life Sciences Research Foundation. This research was also funded by NIH grant K99AI173531 to D.W.M and AI122528 to B.A.G.

---

**Figure S1.**
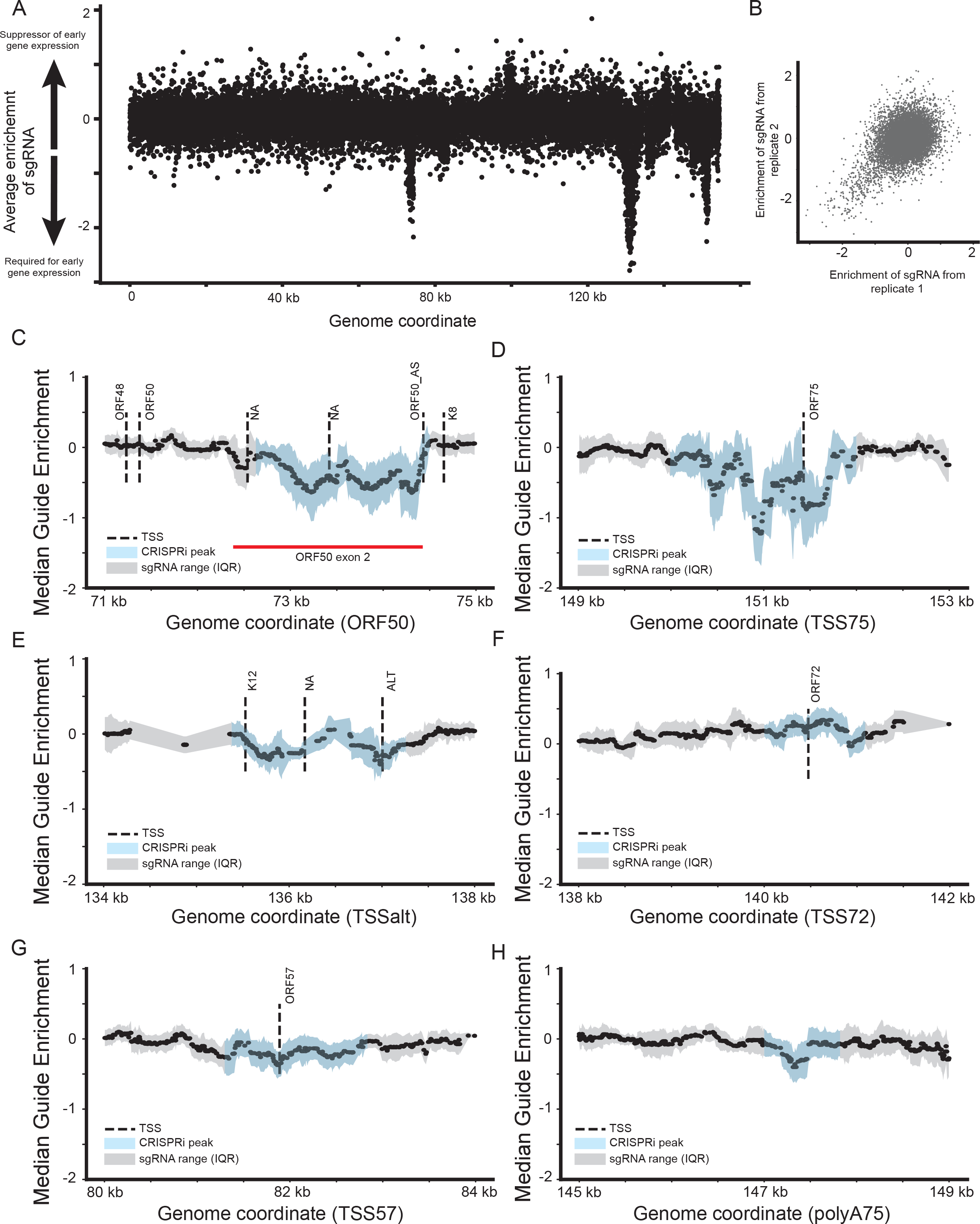
*Supplementary screen data.* A) Enrichment of individual guides at the ORF68 locus. Each dot represents a single guide, with the target location displayed on the x-axis and the average enrichment from two replicates on the y-axis. B) Reproducibility of guide enrichments from two replicates. C-H) Smoothed enrichment of guides at indicated locus with annotated transcription start sites^35^. For each guide, the median enrichment of a 100 bp window centered at the target locus was calculated along with an interquartile range (IQR) to represent the range of values. Median value is shown as a point and IQR is shown as shaded region. Regions significant (p<10^-11^) in sliding window analysis are shown in blue. NA indicates unannotated TSSs.

**Figure S2.**
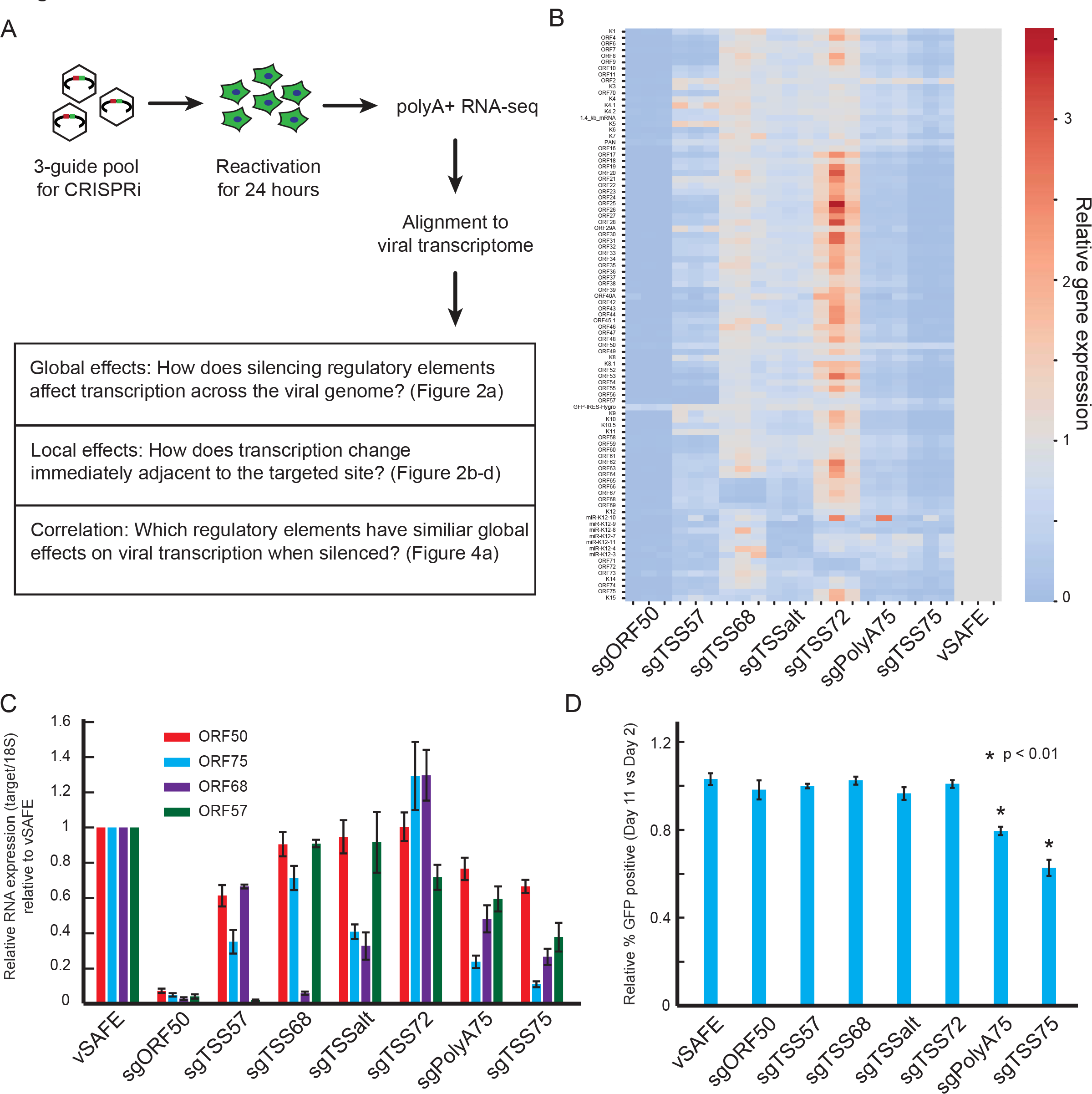
*Supplementary for RNA-seq*. A) Schematic showing set-up of RNA-seq experiment on CRISPRi cells infected with a three-guide pool, reactivated, and polyA+ RNA-seq at 24 hours post-reactivation. B) Heatmap indicating viral gene expression relative to the matched vSAFE replicate of each individual replicate. Rows are presented in genome order. Replicates from three independent reactivations. C) RT-qPCR was used to measure how CRISPRi-based repression of the individual elements indicated on the x-axis influenced the levels of ORFs 50, 75, 68 and 57 mRNA. Error bars represent standard error of four independent reactivations. D) Effect on latency measured by loss of virally encoded EGFP expression over 10 days. Values are presented relative to parental cells, and error bars are standard error from three parallel replicates. P-values are calculated by t-test.

**Figure S3.**
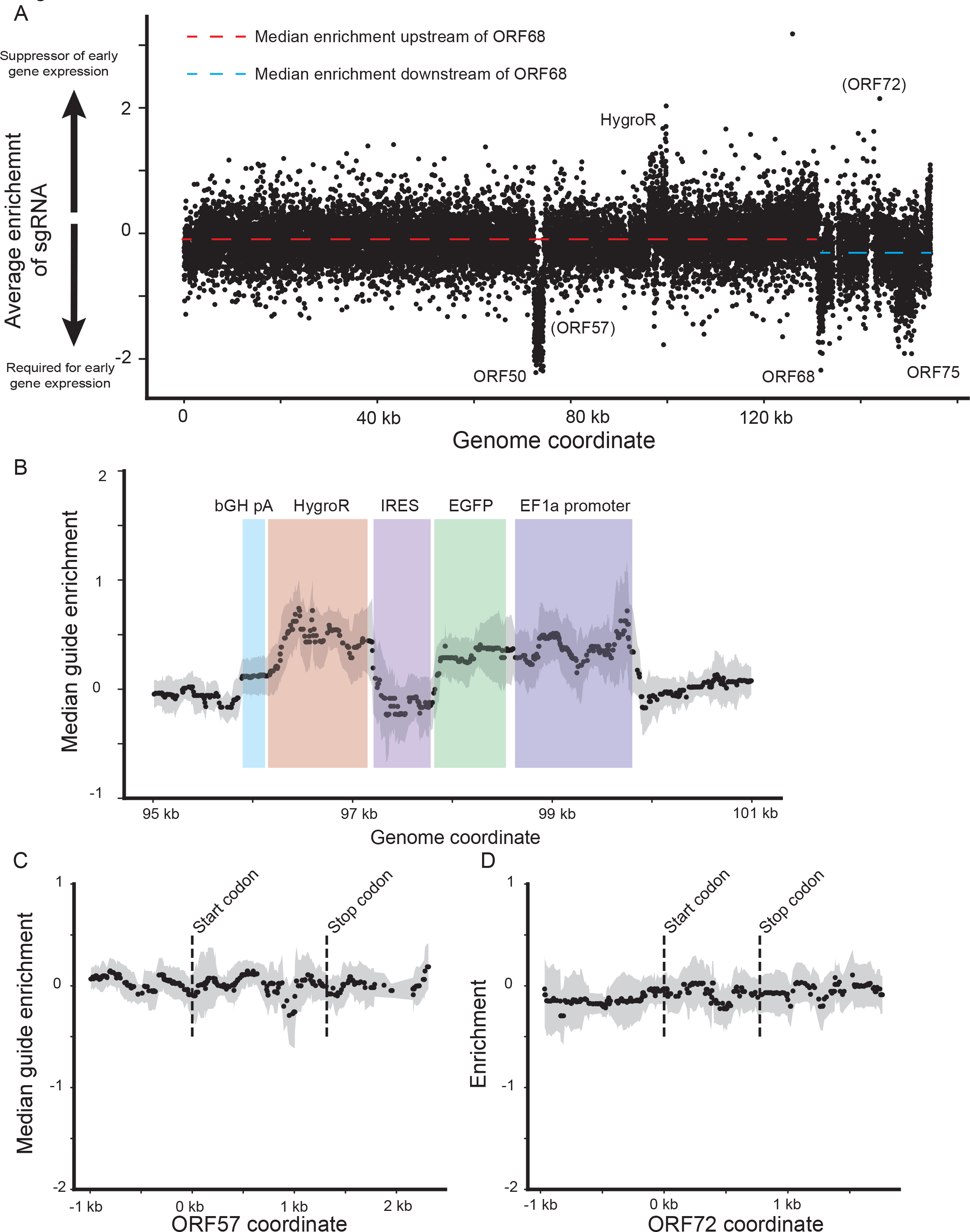
*Coding screen supplement*. A) Enrichment of individual guides. Each dot represents a single guide, with the target location displayed on the x-axis and the average enrichment from two replicates on the y-axis. Red and blue dotted lines represent the median guide enrichment for the two regions indicated. B-D) Median smoothed data from coding screen at indicated locus. For each guide, the median enrichment of a 500 bp window centered at the target locus was calculated along with an IQR. Median value is shown as a point and IQR is shown as shaded region. B) For the EGFP-HygroR locus, location of functional units is shown in color. C,D) Exon boundaries are marked by a dotted line.

**Figure S4.**
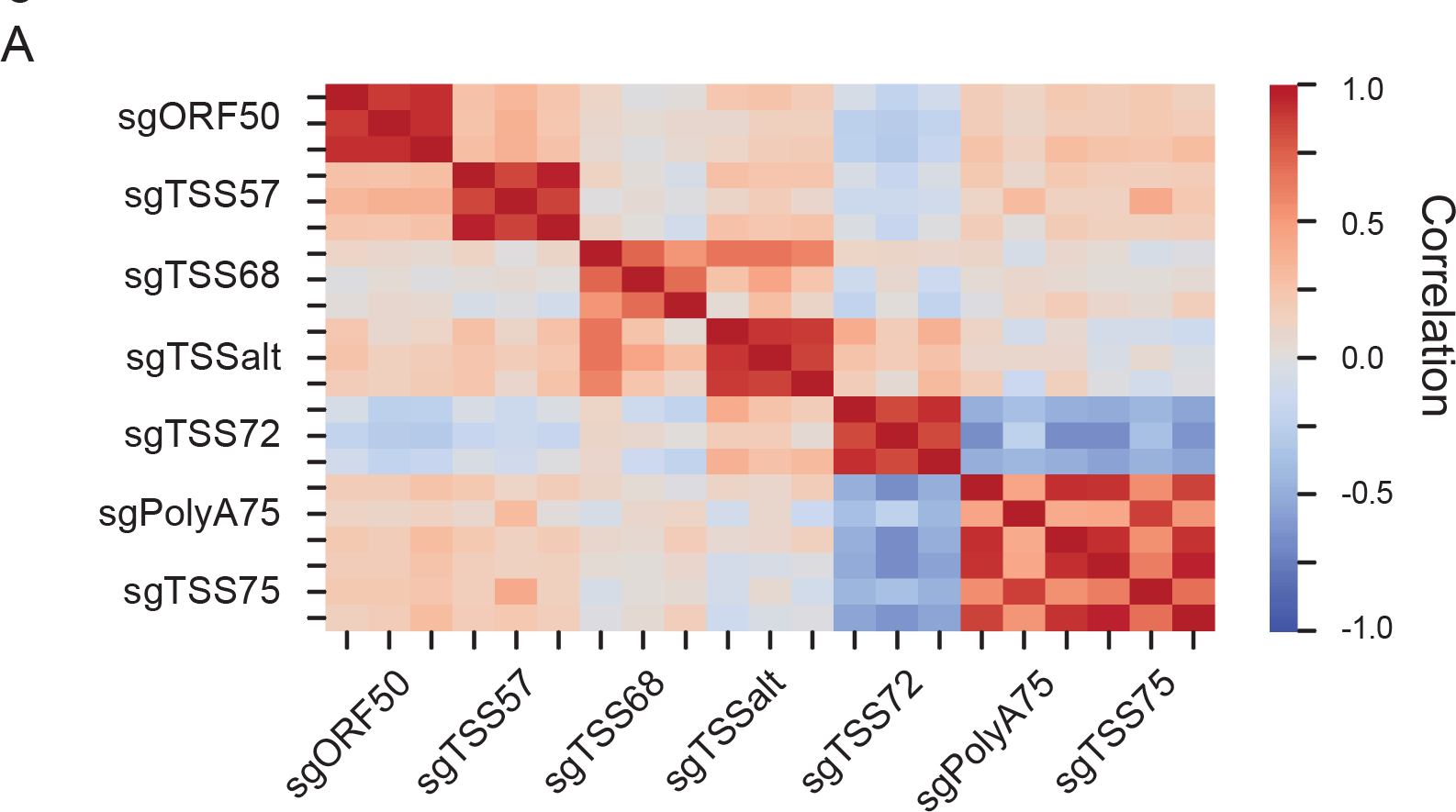
*Supplementary mapping data.* A) Individual replicate correlation among RNA-seq.

**Figure S5.**
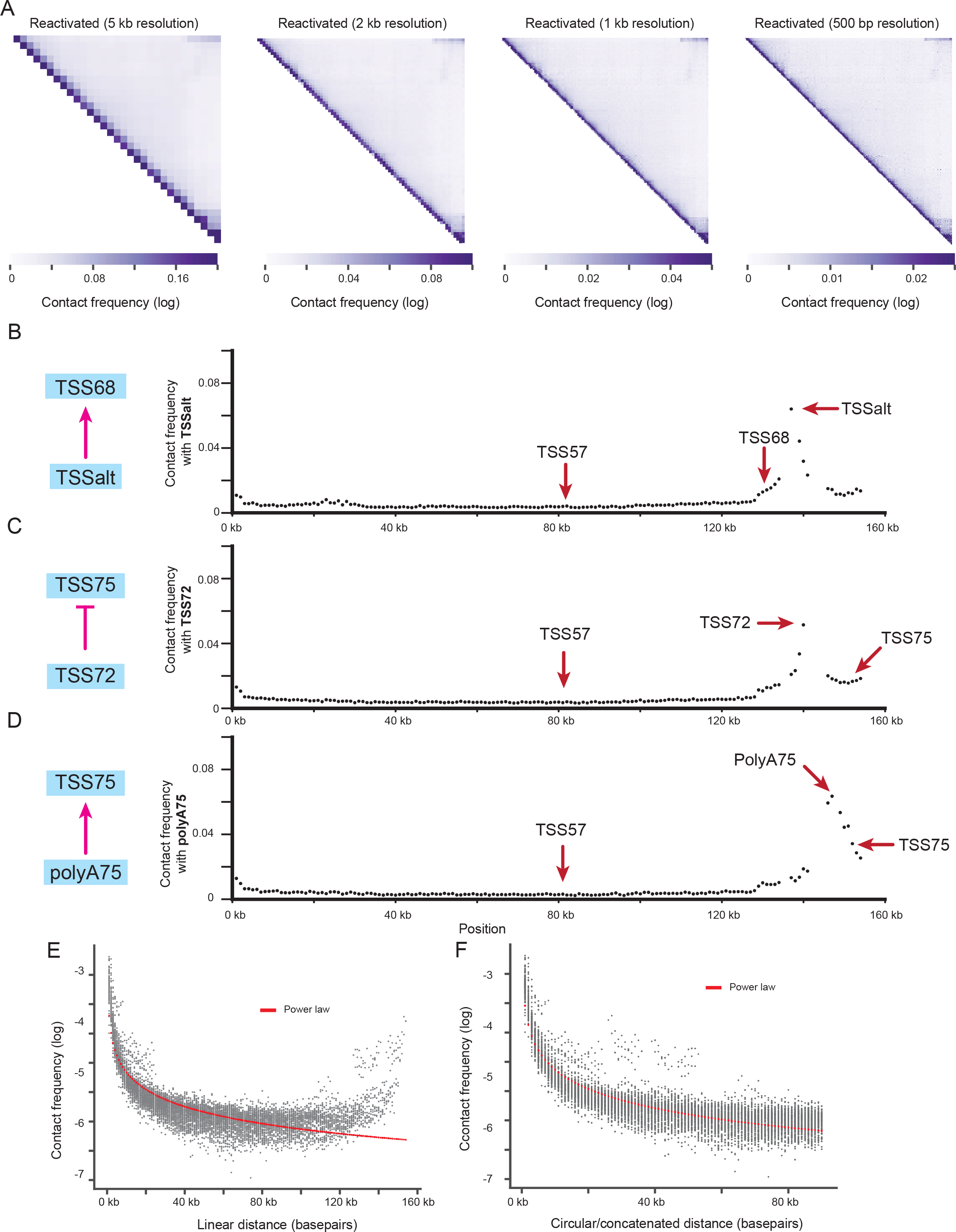
*Hi-C data supplement.* A) Contact frequency of reactivated sample at 5 kb, 2 kb, 1 kb, and 500 bp resolution. B-D) Contact frequency between B) TSSalt, C) TSS72, and D) polyA75 and other locations in the viral genome at 1 kb resolution. E,F) Observed relationship between observed contact frequency and distance between regions when calculated using E) a linear distance metric or F) a circular/concatenated distance metric.

**Figure S6.**
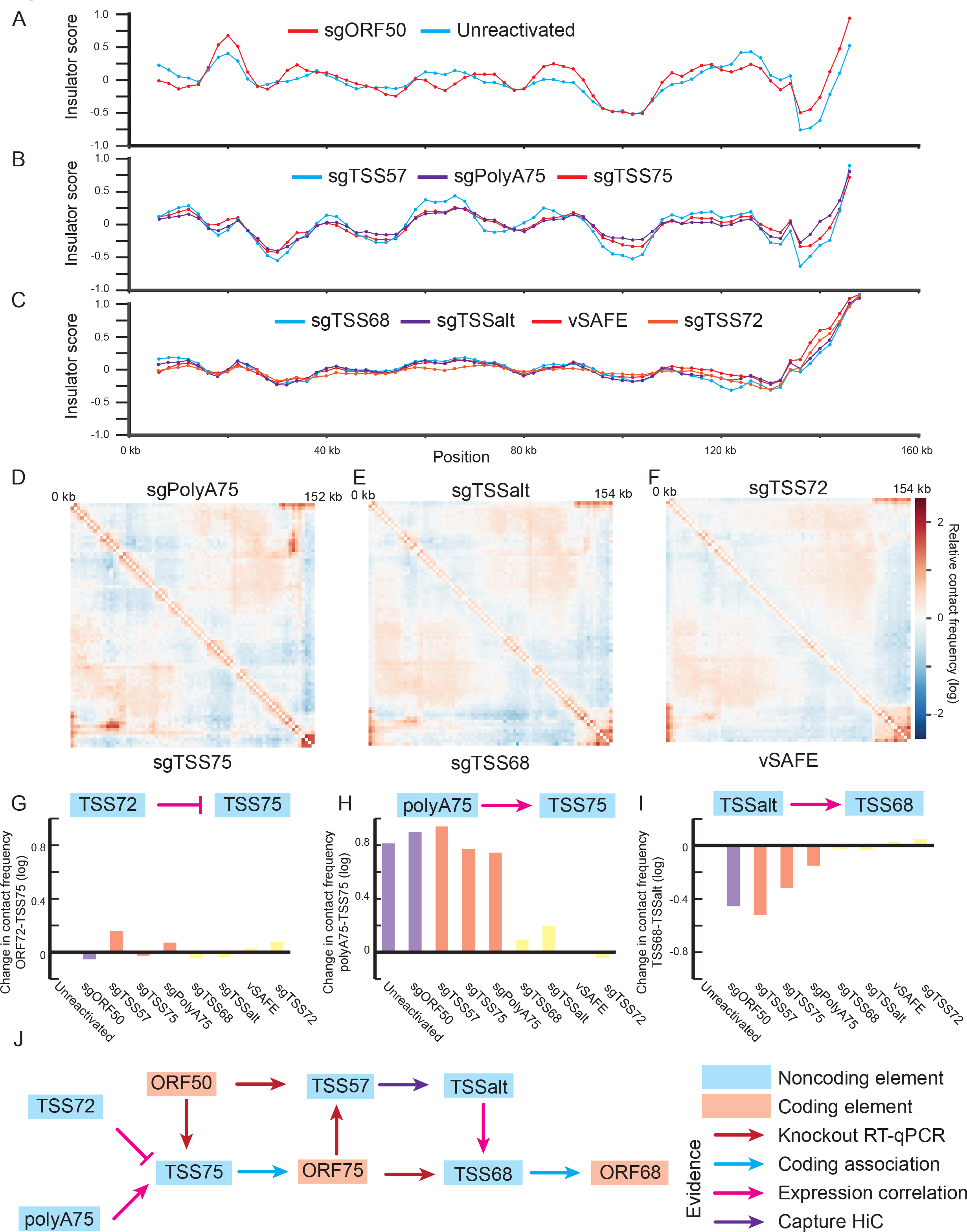
*Structural changes supplement*. A-C) Insulator scores for marked conditions measuring the relative frequency of reads crossing a given location. A more negative value indicates a strong insulator. Local minima were used to define the regions marked in Figure 6e-g. D-F) Relative contact frequency at 2 kb resolution for D) sgPolyA75 and sgTSS75, E) sgTSSalt and sgTSS68, and F) sgTSS72 and vSAFE samples. Positive values indicate greater interaction than expected. G-I) Change in contact frequency from reactivated cells between G) TSS72 and TSS75, H) polyA75 and TSS75, and I) TSS68 and TSSalt at 2 kb resolution. J) Final model for regulatory circuit based on all available data.

## Notes

### Competing Interest Statement

The authors have declared no competing interest.

### Summary of Updates

This revision includes significant additional data encompassing viral capture Hi-C, which is now included as figures 5 and 6, along with supplemental figures 5 and 6 and supplementary data 7. Title, abstract, and text have been updated appropriately.

